# Structure-guided deimmunisation of Fel d 1 suppresses allergic effector functions ex vivo

**DOI:** 10.64898/2026.03.18.712552

**Authors:** Kyungsoo Ha, Soo-Young Yum, Hyeji Kwon, Ho Won Lee, Okjae Koo, Kyeong-Hyeon Eom, Jungjoon K. Lee, Yoonjoo Choi

## Abstract

Allergic responses to the major cat allergen Fel d 1 are primarily driven by immunoglobulin E (IgE) cross-linking on effector cells. While conventional allergen-specific immunotherapies often rely on whole-allergen extracts with inherent risks of systemic reactions, rationally engineered variants offer a safer path to desensitisation. We present a structure-guided strategy to deimmunise Fel d 1 by selectively disrupting crucial IgE-binding interfaces. Using a computational pipeline integrating structural mapping and immunoinformatics, we designed 30 single-point mutations targeting immunodominant regions. Ex vivo functional evaluation using sera from cat-allergic individuals demonstrated that specific variants, notably K29G in Chain 1, exhibited significantly diminished IgE reactivity and profoundly suppressed basophil activation. Furthermore, CRISPR/Cas9-mediated introduction of the K29G mutation into feline fibroblasts confirmed that the substitution preserves fundamental cellular proliferation. This work demonstrates that targeted computational design can not only yield superior candidates for safer immunotherapeutics but also establish a viable foundation for generating genetically edited, hypoallergenic cats.

## Introduction

Allergens are substances capable of triggering hypersensitive immune responses in susceptible individuals^1,2^. They can originate from a wide range of sources from foods and pollens to animal dander^3^, which make allergic diseases a significant health concern^4^. Among animal-derived allergens, cat allergy is one of the most common pet allergies, affecting roughly 20% of the global population^5^. One of the major allergens in cat allergy is Fel d 1, a protein found in cat saliva, skin, and fur, which accounts for about 90% of allergic reactions to cats^6^. Fel d 1 is a small secreted heterodimeric glycoprotein (∼35-40 kDa) composed of two polypeptide chains (chain 1 and chain 2 of ∼18 kDa each)^6,7^. Its precise biological function remains largely unclear^8^, but clinically Fel d 1 is among the most significant indoor allergens. Indeed, 90-95% of individuals with cat allergy have IgE antibodies against Fel d 1^9^ and many of them suffer from feline-induced atopy due to the allergen^10^. Moreover, this robust IgE response is inherently T cell-dependent requiring allergen-specific CD4+ T-helper cells to initiate and maintain the allergic cascade^11^.

Managing cat allergy is challenging, as avoiding exposure to pets or ubiquitous allergen reservoirs is often impractical. Symptomatic treatments (e.g., antihistamines, inhaled corticosteroids) provide relief but do not address the underlying IgE-mediated sensitivity. Currently, allergen-specific immunotherapy (AIT), which involves administering increasing doses of cat allergen extract to build tolerance, is the sole disease-modifying treatment available^12^. Conventional immunotherapy, however, is a prolonged and burdensome process. It may cause severe allergic reactions since it utilises the native allergen^13^. Moreover, its efficacy can vary greatly between patients^14^.

These limitations have spurred the exploration of alternative strategies to neutralise Fel d 1 at the source or develop safer therapeutics. Attempts to decrease allergen shedding via nutritional interventions, such as feeding cats egg yolk powder containing anti-Fel d 1 IgY antibodies, offer limited mitigation^15^. Similarly, efforts to develop altered peptide immunotherapies^16^ aimed at inducing tolerance without provoking IgE cross-linking have shown limited success, with early candidates failing to demonstrate significant clinical benefit in Phase III trials^17,18^. Given the protein nature of Fel d 1, another concept has been to create hypoallergenic variants of the molecule with mutations that reduce IgE reactivity while preserving enough structure to induce immunological tolerance. This approach, called “hypoallergen” design, has been facilitated by knowledge of Fel d 1 structure and the mapping of key IgE-binding regions^19^. Indeed, the crystal structure of Fel d 1 revealed distinct surface regions responsible for IgE binding. However, to date, no Fel d 1 variants has progressed to clinical use, and hypoallergen design efforts have mainly focused on IgE epitopes alone. Targeting T-cell epitopes is also equally important for allergenicity^20,21^, and thus removing T-cell epitopes can be an effective strategy to reduce allergenicity.

Concurrently, modern gene-editing technologies have opened a direct route to produce hypoallergenic cats. Recent CRISPR/Cas9 studies have provided proof-of-concept that the Fel d 1 gene may be knocked out^22,23^. However, the long-term biological consequences of completely ablating Fel d 1 remain uncertain. Therefore, retaining Fel d 1 expression while rationally redesigning its structure to eliminate allergenicity may represent a promising, sustainable approach. Engineering cats to produce a deimmunised Fel d 1 variant would theoretically preserve its unknown physiological functions while directly addressing the clinical root of the allergy.

To resolve these challenges, we present a structure-guided strategy to deimmunise Fel d 1. Leveraging recent advances in computational biology and protein design algorithms^24,25,26^ alongside well-characterised IgE and T-cell epitope maps, we generated novel variants of Fel d 1. We hypothesised that targeted single-point mutations within structural hotspots could disrupt critical IgE-binding interfaces without destabilising the protein^27^. Using a computational pipeline, we designed these variants and evaluated their efficacy in vitro and ex vivo. To establish translational feasibility, we employed CRISPR/Cas9 genome editing to introduce the most successful mutation into feline fibroblasts in culture. We found that the edited cells proliferate normally, demonstrating no observable detrimental effects on fundamental cellular fitness. These results show how modern protein design and gene editing technologies can converge to address clinical challenges in allergen avoidance and therapy.

## Results

### Structure-guided computational design of deimmunised Fel d 1 variants

To rationally design hypoallergenic variants of Fel d 1, we first established a baseline recombinant construct, rFel d 1, comprising a single-chain fusion of the native heterodimeric chain 1 and chain 2 (**Supplementary Table S1**). The three-dimensional structure of Fel d 1 (PDB ID: 1ZKR, chain A) was used as the structural template for our computational deimmunisation strategy (**Fig. 1A**). To ensure that targeted mutations would disrupt immune recognition without destabilising the protein core, we quantified the solvent accessibility of each amino acid using the Define Secondary Structure of Proteins (DSSP) program^28,29^. Only residues with a solvent accessibility score greater than 20 were classified as surface-exposed and considered viable candidates for mutagenesis (**Fig. 1B**).

**Figure 1.**
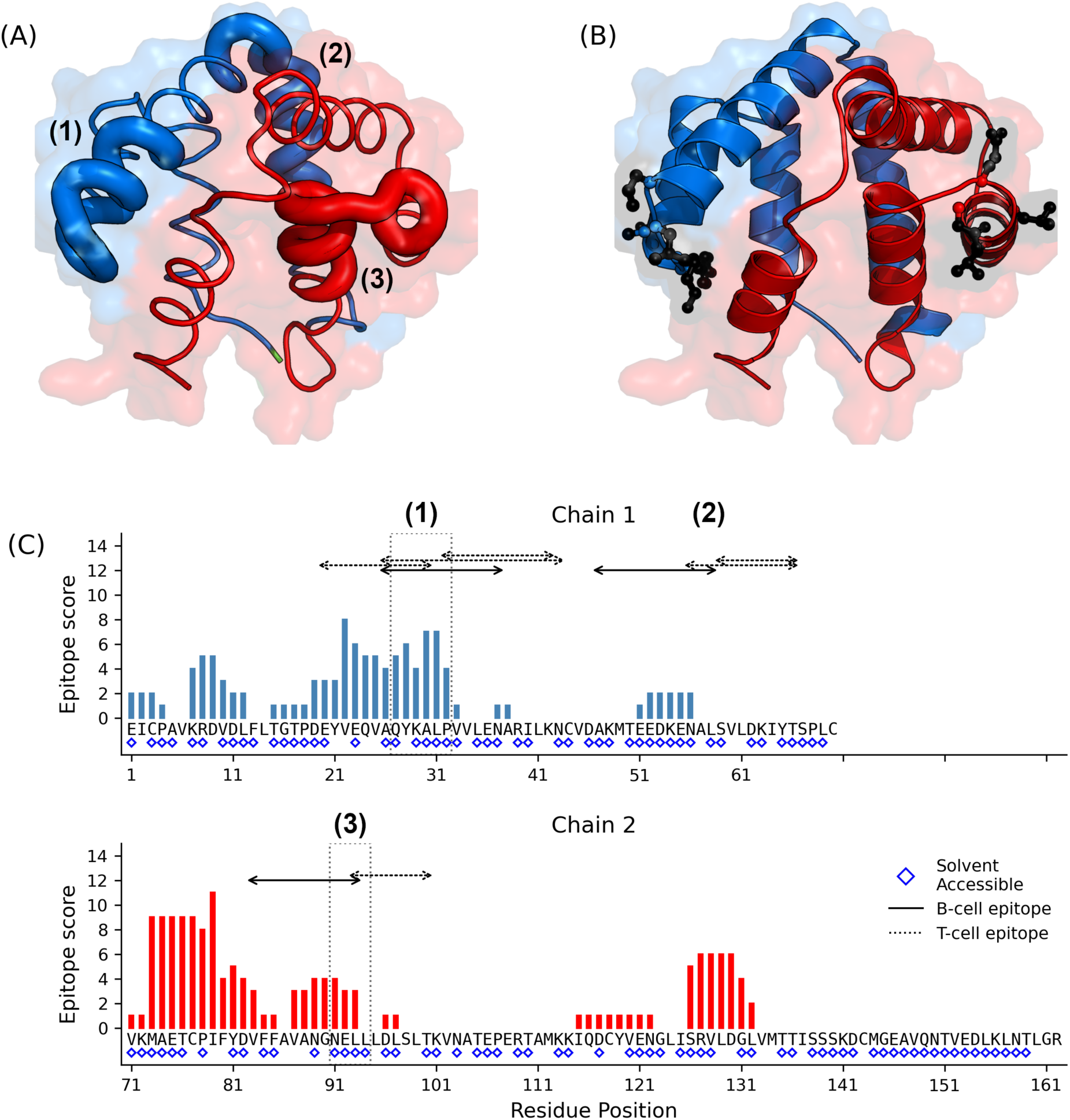
Structural and immunological rationale for Fel d 1 variant design. (A) Surface representation of the Fel d 1 structure highlighting three known B-cell epitopes. (B) Location of designed mutation sites (black sticks) targeting epitope-rich regions across both chains. (C) Predicted MHC class II binding affinity across the Fel d 1 sequence using the IEDB-recommended algorithm. Overlay of B-cell and T-cell epitopes illustrating a high degree of overlap in epitope 1.

Previous epitope mapping studies^30,31,32,33^ have identified three major IgE-binding surfaces, designated B-cell epitopes 1, 2, and 3, distributed across the protein structure (The known major T- and B-cell epitopes are listed in **Supplementary Table S2**). While conventional hypoallergen design often focuses exclusively on these B-cell footprints, we sought a more comprehensive structural redesign. Therefore, we computationally mapped the density of predicted CD4+ T-cell epitopes across the nFel d 1 sequence using the IEDB-recommended pMHC-II binding prediction method^34,35,36^. To ensure maximal population coverage, we queried the full reference set of 27 common human leukocyte antigen (HLA) class II alleles^37^.

This immunoinformatic analysis revealed numerous high-affinity MHC-II binding predictions clustering near the known B-cell epitopes (**Fig. 1C**). This convergence was particularly evident within epitope 1 (residues 25–38) on chain 1, which exhibited the highest degree of structural overlap between predicted T-cell epitopes and surface-exposed IgE contact residues, forming a dual-epitope immunological hotspot. Based on this structural logic, we prioritised epitopes 1 and 3 as the primary regions for targeted disruption.

Initial efforts focused on establishing a reliable recombinant production pipeline for the wild-type rFel d 1 control. The protein was expressed in *E. coli* and subsequently refolded using a rapid detergent-based renaturation protocol. The integrity of this production pipeline was verified via SDS-PAGE analysis (**Supplementary Fig. S1**). The purified rFel d 1 resolved at the expected molecular weight (∼18–20 kDa) and exhibited high purity, comparable to the natural commercial allergen (nFel d 1). The structural fidelity and proper conformational refolding of this recombinant construct were subsequently confirmed by its ability to bind human IgE at levels comparable to nFel d 1 (**Fig. 2B**).

**Figure 2.**
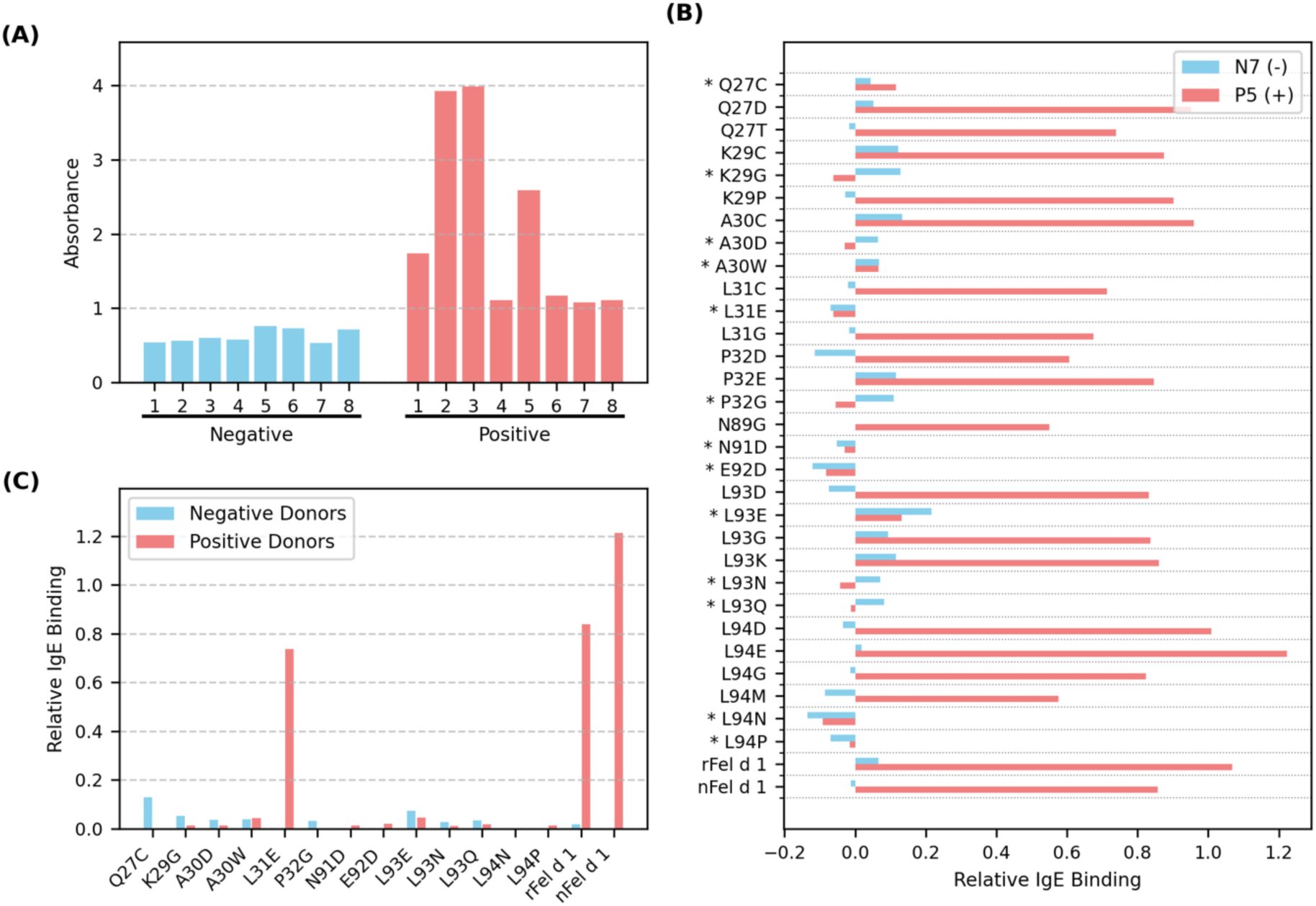
IgE binding activity of wild-type and variant Fel d 1 proteins. (A) ELISA screening of sera from allergic (positive) and non-allergic (negative) donors against natural Fel d 1 (nFel d 1). (B) Initial IgE binding screen of hypoallergenic mutations using sera from a positive donor (P5) and negative donor (N7). (C) Comprehensive screen of selected variants using sera from all IgE-positive donors. While most candidates retained low IgE reactivity, L31E failed to reduce IgE binding in the broader cohort.

We then designed a library of 30 single-point mutations (15 per chain, **Table 1**). Mutations were strictly confined to surface-exposed residues within or immediately adjacent to the identified immunological hotspots, selected for their predicted ability to disrupt pMHC-II binding events while preserving the overall structural architecture. These 30 variants were expressed and refolded applying the identical, validated detergent-based protocol, yielding a robust, soluble library for downstream functional immunological screening.

**Table 1.** Initial IgE-binding screen of 30 Fel d 1 variants. ELISA results from positive (P5) and negative (N7) donor sera for all 30 single-point mutants, wild-type rFel d 1, and nFel d 1.

| Chain | Position (Solvent accessibility) | Variant | N7 (-) | P5 (+) | pMHC-II binding reduction | Chosen for the next round |
| --- | --- | --- | --- | --- | --- | --- |
|  |  | rFel d 1 | 0.5536 | 1.5233 | 0 |  |
|  |  | nFel d 1 | 0.4767 | 1.3122 | 0 |  |
|  |  | Negative | 0.4882 | 0.4551 | - |  |
| 1 | 27 (138) | Q27C | 0.5323 | 0.5707 | 13 | * |
|  |  | Q27D | 0.5393 | 1.4062 | 20 |  |
|  |  | Q27T | 0.4708 | 1.1932 | 8 |  |
|  | 29 (68) | K29C | 0.6096 | 1.3301 | 41 |  |
|  |  | K29G | 0.6163 | 0.3935 | 31 | * |
|  |  | K29P | 0.4599 | 1.3573 | 35 |  |
|  | 30 (65) | A30C | 0.622 | 1.4147 | 29 |  |
|  |  | A30D | 0.5529 | 0.4257 | 34 | * |
|  |  | A30W | 0.5566 | 0.5212 | 20 | * |
|  | 31 (90) | L31C | 0.4679 | 1.1689 | 44 |  |
|  |  | L31E | 0.4177 | 0.3932 | 37 | * |
|  |  | L31G | 0.4709 | 1.1303 | 40 |  |
|  | 32 (100) | P32D | 0.3742 | 1.0607 | 11 |  |
|  |  | P32E | 0.6036 | 1.3007 | 14 |  |
|  |  | P32G | 0.5972 | 0.3992 | 18 | * |
| 2 | 89 (56) | N89G | 0.4884 | 1.0059 | 7 |  |
|  | 91 (74) | N91D | 0.4362 | 0.4257 | 8 | * |
|  | 92 (104) | E92D | 0.368 | 0.3733 | 6 | * |
|  | 93 (129) | L93D | 0.4135 | 1.2868 | 43 |  |
|  |  | L93E | 0.7052 | 0.5874 | 40 | * |
|  |  | L93G | 0.5808 | 1.2918 | 45 |  |
|  |  | L93K | 0.6041 | 1.3158 | 36 |  |
|  |  | L93N | 0.5595 | 0.4124 | 44 | * |
|  |  | L93Q | 0.5705 | 0.4428 | 44 | * |
|  | 94 (94) | L94D | 0.4538 | 1.4634 | 16 |  |
|  |  | L94E | 0.5069 | 1.6789 | 16 |  |
|  |  | L94G | 0.475 | 1.2786 | 19 |  |
|  |  | L94M | 0.402 | 1.0305 | 7 |  |
|  |  | L94N | 0.3532 | 0.3633 | 16 | * |
|  |  | L94P | 0.4189 | 0.4403 | 24 | * |

### Identification of hypoallergenic variants through IgE binding screens

To identify Fel d 1 variants with reduced IgE reactivity, We first quantified IgE binding to natural Fel d 1 (nFel d 1) across all donor sera to establish baseline allergen-specific responses and identify individuals with robust IgE titres (**Fig. 2A**). Based on this reactivity profile, we selected one highly reactive IgE-positive donor (P5) and one negative donor (N7) to perform a preliminary screen of the entire 30-variant library (**Fig. 2B**). As a control, recombinant wild-type Fel d 1 (rFel d 1) was also included in the assay to validate the allergenicity of our refolded protein. rFel d 1 displayed IgE-binding levels that were comparable to those of nFel d 1 in the P5 serum.

Within this preliminary assay, we evaluated the IgE-binding capacity of each single-point mutation. Several variants, particularly those targeting the dual-epitope hotspot within epitope 1, exhibited a profound reduction in IgE signal in the P5 serum, while maintaining negligible background binding in the negative control (**Table 1**).

To rigorously validate these findings across broader immunological profiles, we advanced the most promising candidates to a comprehensive IgE binding assay utilising sera from the complete cohort of IgE-positive donors (**Fig. 2C**). While several engineered variants successfully maintained their reduced IgE reactivity profiles across multiple donors, this extended screen also exposed critical inter-individual variability. For instance, the L31E variant, which initially appeared highly effective in the P5 screen, failed to consistently evade IgE binding across the broader cohort (**Supplementary Table S3**). The screening process successfully isolated a subset of Fel d 1 variants that consistently evaded IgE recognition, establishing a validated foundation for subsequent functional evaluation in cellular assays.

### Lead candidate demonstrates reduced basophil activation

To determine whether the observed reductions in IgE binding translated into functional hypoallergenicity, we performed a basophil activation test (BAT). In this assay, peripheral blood basophils are sensitised with serum IgE from allergic donors and subsequently challenged with candidate allergens. Basophil activation is quantified by flow cytometric detection of surface markers such as CD63 and CD123 that provide a physiologically relevant measure of effector cell degranulation.

Initial validation of cellular responsiveness and assay integrity was conducted using the non-specific stimulant fMLP, employing basophils sensitised with serum from the highly reactive donor P5 (**Fig. 3A** and **Supplementary Fig. S2**). Having established these assay parameters, we evaluated the lead hypoallergenic variant (K29G) utilising a distinct cohort of Fel d 1-reactive donors: P2, P3, and P8. Donors P2 and P3 were specifically selected as they exhibited the most profound IgE binding responses in the preceding ELISA screens (**Fig. 2A**). While donor P8 displayed a more moderate IgE reactivity profile, this sample was strategically included due to its high primary basophil yield. Basophils sensitised with sera from each of these three donors were stimulated with either wild-type recombinant Fel d 1 (rFel d 1) or the engineered K29G variant.

**Figure 3.**
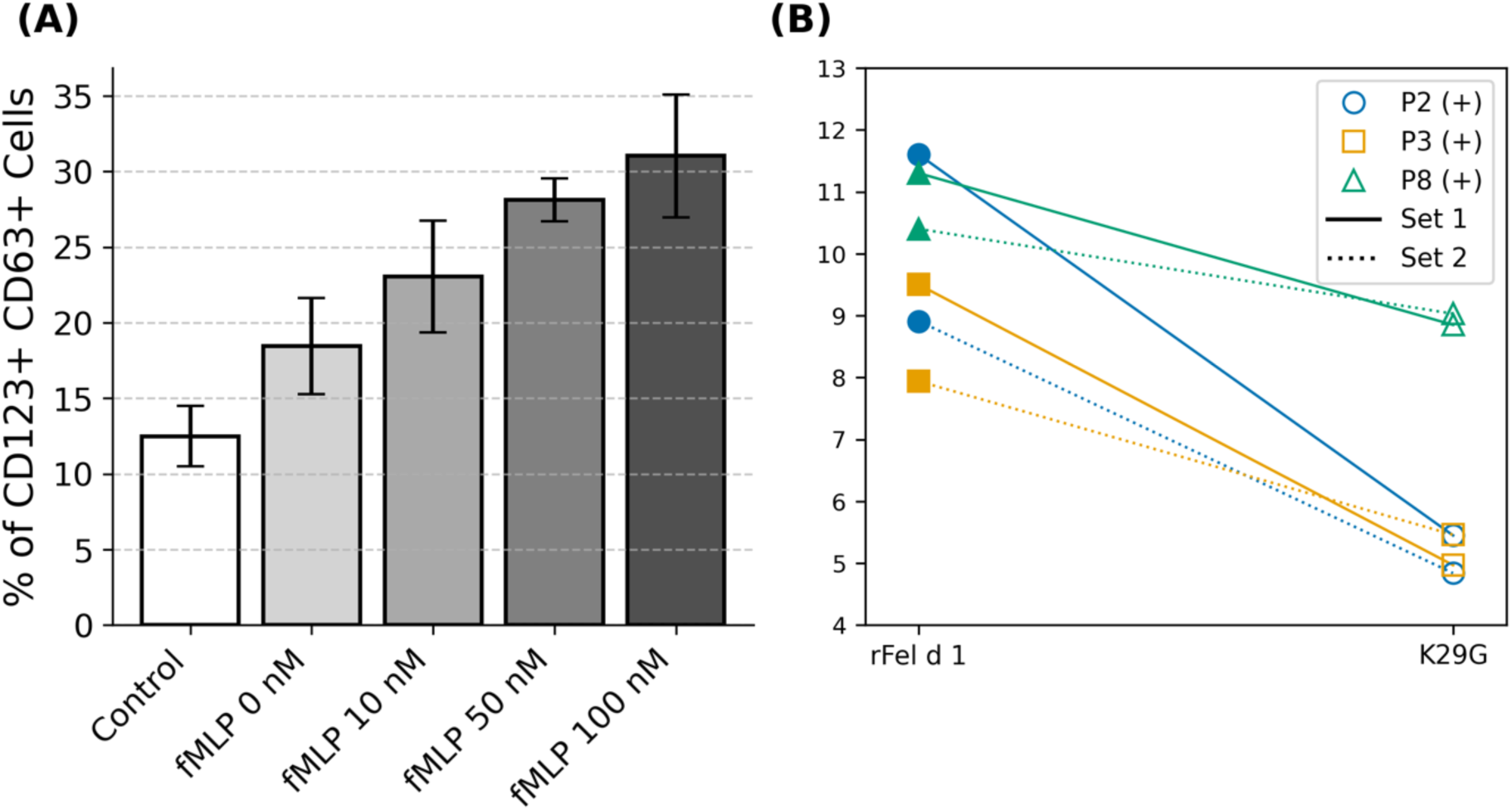
Basophil activation assays confirm functional hypoallergenicity. (A) Dose-dependent activation of P5 basophils by fMLP validates assay responsiveness. (B) Stimulation of basophils from donors P2, P3, and P8 with rFel d 1 or variant K29G (duplicated). Wild-type protein induced strong CD63+ activation, while K29G elicited significantly reduced responses.

As expected, rFel d 1 induced pronounced basophil activation in all donors, consistent with its strong IgE-binding capacity. In contrast, stimulation with the K29G variant resulted in a markedly attenuated activation response, with CD123+ CD63+ basophil frequencies reduced to near-background levels (**Fig. 3B**). The suppression of activation was consistent across all three donors and confirmed by the detailed FACS analyses (**Supplementary Fig. S3**). These results provide compelling functional evidence that the engineered mutations not only diminish IgE recognition but also effectively abrogate downstream basophil activation.

### Proliferation analysis of feline cell lines

Translational feasibility of mitigating allergen exposure at the source necessitates native expression of the engineered variant. We performed CRISPR/Cas9-mediated genome editing on feline fetal fibroblasts using a ribonucleoprotein (RNP) complex paired with a single-stranded oligodeoxynucleotide (ssODN) donor template. Following transfection, we successfully established cell lines harbouring the site-specific K29G substitution, which was subsequently confirmed via Sanger sequencing (**Supplementary Fig. S4**).

A critical prerequisite for in vivo gene-editing application is ensuring that the introduced mutation does not disrupt intrinsic cellular machinery. We longitudinally monitored the population doubling time (PDT) of the edited cell line compared to wild-type (WT) fibroblasts from passages 8 to 19, determining whether the K29G substitution impacted fundamental cellular fitness. Both the WT and K29G lines exhibited characteristic phenotypic variability, with PDT values generally fluctuating between 10 and 40 hours (**Fig. 4**). While transient proliferation delays were observed, specifically at passage 9 for the WT (∼48 h) and passage 12 for the K29G-edited line (∼67 h), both cultures demonstrated a clear trend towards stabilisation in subsequent passages.

**Figure 4.**
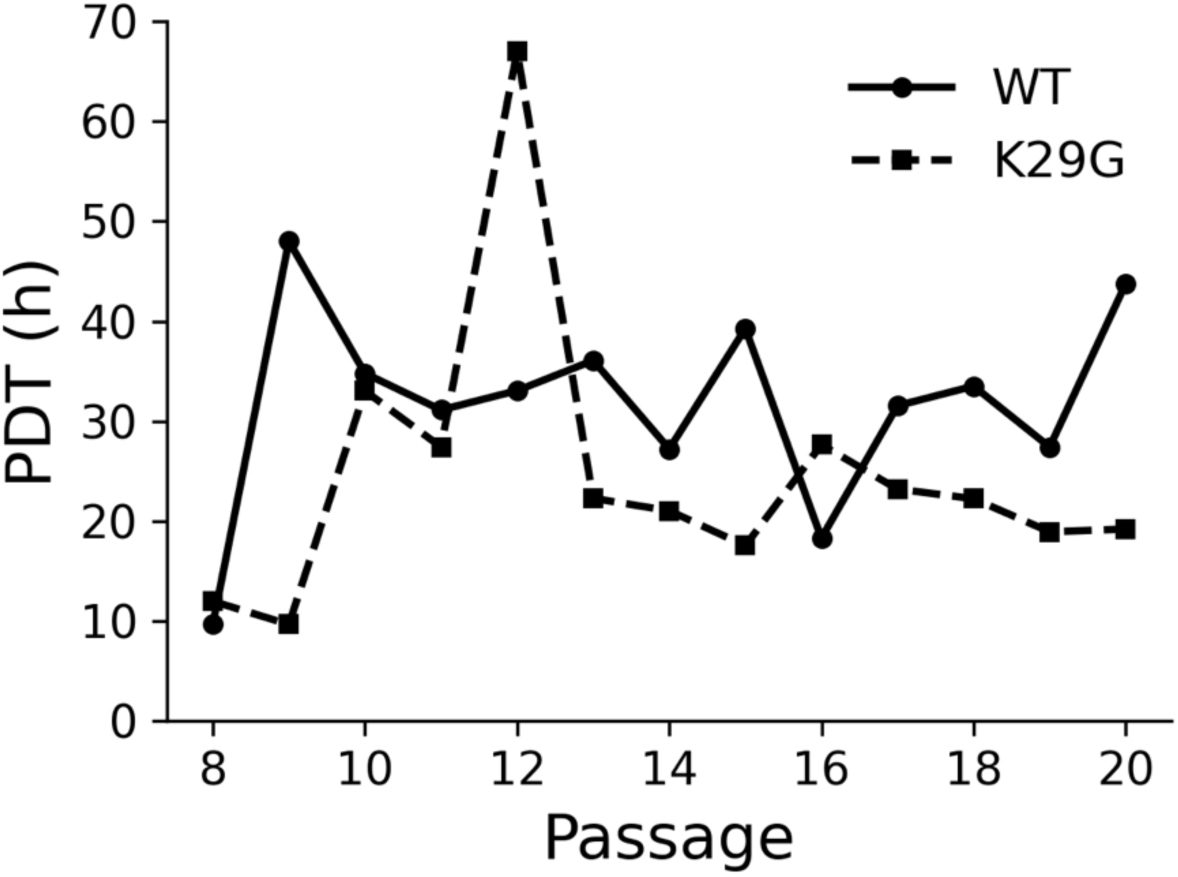
Proliferation analysis of wild-type (WT) and K29G-edited feline fetal fibroblasts. Population doubling time (PDT) monitoring (P8-19) reflected inherent variability of primary cell lines. Subsequent stabilization indicates that the K29G substitution does not compromise fundamental cellular fitness.

Crucially, we detected no sustained growth inhibition or overt morphological abnormalities in the edited cells over the extended monitoring period. The overall proliferative capacity of the K29G fibroblasts remained consistent with that of the WT controls throughout the longitudinal monitoring phase. These results indicate that the targeted K29G substitution does not compromise the fundamental cellular processes of feline fibroblasts. Consequently, this data provides a vital in vitro proof-of-concept, supporting the biological viability of utilising structural deimmunisation paired with gene-editing technologies for sustainable allergen avoidance.

## Discussion

Therapeutics for feline-induced atopy present significant practical difficulties due to severe immunogenicity. In this study, we demonstrated the successful design, production, and functional validation of hypoallergenic Fel d 1 variants using a rational, dual-epitope targeting strategy. By focusing on regions of overlap between known B-cell epitopes and predicted MHC class II T-cell epitopes, we aimed to simultaneously disrupt IgE recognition and computationally minimise pMHC-II binding potential. Epitope 1, in particular, emerged as a prominent immunological hotspot, showing dense overlap between surface-exposed IgE contact residues and high-affinity MHC-II peptide predictions. Guided by this dual-epitope logic, we generated 30 single-point mutations distributed across both chains of Fel d 1 and expressed each variant as a soluble recombinant protein.

Our recombinant wild-type Fel d 1 (rFel d 1) was successfully expressed, refolded, and validated to maintain comparable IgE-binding activity to native Fel d 1, serving as a reliable internal control. This robust baseline allowed us to proceed with screening against human sera. We employed the basophil activation test (BAT) as a functional assay to measure the cellular effects of a designed variant in epitope 1 (K29G). We challenged sensitised basophils from high-titre donors. Wild-type rFel d 1 induced robust activation, whereas the K29G variant triggered a reduction in basophil activation. This direct suppression of basophil degranulation provides strong evidence that our variants are functionally hypoallergenic and capable of dampening the effector phase of the allergic response.

Beyond its potential as injectable immunotherapies, with recent advances in CRISPR/Cas9 genome editing technology, it is now conceivable to introduce validated hypoallergenic mutations directly into the feline genome. Gene-edited cat cells that naturally express a low-allergenicity version of Fel d 1 could offer a transformative approach to eliminating the source of sensitisation altogether. Such a strategy would not only benefit allergic individuals but also reduce the need for lifestyle compromises for cat owners worldwide. This study thus marks a critical step toward both immunotherapeutic and genomic solutions for cat allergy.

## Methods

### Structural and immunoinformatic analysis

The crystallographic structure of Fel d 1 (PDB ID: 1ZKR, chain A) was retrieved from the Protein Data Bank. Solvent accessibility was calculated employing the Define Secondary Structure of Proteins (DSSP) program^28,29^. Residues exhibiting a solvent accessibility score strictly greater than 20 were classified as surface-exposed. Structural visualisations and epitope mapping renderings were generated utilising PyMOL (version 3.0).

Prediction of pMHC-II binding was executed using the IEDB-recommended method (version 2.18) against a full reference set of 27 human HLA class II alleles to maximise population coverage^37^. A binding event was assigned a binary score of 1 if a peptide was predicted to be a binder (percentile rank < 10%) for a specific allele, and 0 otherwise. Binding scores for each 15-mer peptide were calculated by summing the total binding events across all 27 alleles, yielding a maximum potential score of 27. This cumulative score facilitated the mapping of predicted T-cell epitope density onto the protein structure and the assessment of spatial proximity to established B-cell epitopes and targeted mutation sites.

### Human cohort recruitment and sample collection

All protocols for the recruitment of human subjects and the collection of biological materials were approved by the Institutional Review Board (IRB) of the Korea Advanced Institute of Science and Technology (KAIST; approval number KH2019-74). Thirty volunteers were initially recruited at the KAIST clinic based on self-reported clinical histories of cat allergy. Following a preliminary screening using a commercial cat allergy self-test (Imutest Cat Allergy Kit), 16 donors were selected for subsequent immunological assays. This cohort comprised eight positively identified cat-allergic individuals and eight non-allergic controls.

Peripheral blood (30 ml per donor) was drawn for the isolation of serum and peripheral blood mononuclear cells (PBMCs). Serum was immediately separated via centrifugation at 1,000×g for 10 minutes at room temperature, aliquoted, and stored at -80 °C. PBMCs were isolated using standard density-gradient centrifugation. Briefly, blood samples were diluted 1:1 with sterile phosphate-buffered saline (PBS) and layered onto a Ficoll-Paque density gradient medium (GE Healthcare). Following centrifugation at 400×g for 30 minutes at room temperature without braking, the PBMC layer at the plasma-Ficoll interface was carefully collected. Cells were washed twice with PBS via centrifugation at 300×g for 10 minutes at 4 °C, and then resuspended in RPMI 1640 medium supplemented with 10% foetal bovine serum (FBS) for immediate utilisation in basophil activation tests.

### Recombinant protein expression and refolding

A synthetic expression cassette encoding chain 1 and chain 2 fused, incorporating an N-terminal hexahistidine tag, was synthesised (Twist Bioscience) and cloned into a kanamycin-resistant pET-based vector. Chemically competent *E. coli* BL21(DE3) cells (Thermo Fisher Scientific) were transformed and cultured in Luria-Bertani (LB) medium containing 50 µg/ml kanamycin at 37 °C. Upon reaching an optical density (OD600) of approximately 0.4, protein expression was induced using 0.5 mM isopropyl β-D-1-thiogalactopyranoside (IPTG) for 3 hours. Cells were subsequently harvested via centrifugation at 5000×g for 10 minutes at 4 °C.

As recombinant Fel d 1 predominantly accumulates in inclusion bodies, a detergent-assisted solubilisation strategy was employed. Cell pellets were chemically lysed in an ice-cold STE-based buffer supplemented with lysozyme and a protease inhibitor cocktail (Tocris Bioscience). Inclusion bodies were solubilised using a proprietary Detergent Solubilisation Solution (Rapid GST Inclusion Body Solubilisation and Renaturation Kit, Cell Biolabs) for 1 hour on ice. Following centrifugation, the clarified supernatant was treated with a matched Detergent Neutralisation Solution for an additional 1 hour on ice. This procedure facilitated the rapid refolding of the protein into its native conformation without requiring extensive dialysis or redox chemicals.

The neutralised preparation was concentrated using Amicon Ultra centrifugal filter units (Merck MilliporeSigma) and subjected to immobilised metal affinity chromatography. His-tagged Fel d 1 was purified using Ni-NTA spin columns (Thermo Fisher Scientific or Qiagen), washed with PBS containing 25 mM imidazole, and eluted with 250 mM imidazole. Eluates were immediately buffer-exchanged into physiological PBS using Zeba Spin Desalting Columns (Thermo Fisher Scientific) to recover the desalted protein for downstream functional evaluation.

### IgE-binding and basophil activation assays

Allergen-specific IgE binding was quantified employing an indirect enzyme-linked immunosorbent assay (ELISA). High-binding 96-well microtitre plates were coated with Fel d 1 antigens (4 µg/ml) in carbonate-bicarbonate buffer (pH 9.6) and incubated overnight at 4 °C. Plates were washed with PBS containing 0.05% Tween-20 (PBST) and blocked with 1% bovine serum albumin (BSA) in PBST for 1 hour at room temperature. Human serum samples, diluted 1:2 in PBS, were applied and incubated for 2 hours at 37 °C. Following subsequent washes, bound IgE was detected using a horseradish peroxidase (HRP)-conjugated goat anti-human IgE secondary antibody (1:2000 dilution) during a 1-hour incubation at 37 °C. The colourimetric reaction was developed with 3,3’,5,5’-tetramethylbenzidine (TMB) substrate for 10 minutes, terminated with 2 M H₂SO₄, and absorbance was measured at 450 nm. All assays were performed in duplicate.

Functional hypoallergenicity was evaluated via a flow cytometry-based basophil activation test (BAT). Cryopreserved human peripheral blood basophils were thawed and resuspended at a density of 1.5 × 10⁶ cells per 500 µl of medium. Cells were primed with 0.3 nM IL-3 at 37 °C for 10 minutes prior to stimulation. Basophils were subsequently challenged with either the non-specific stimulant fMLP (10, 50, or 100 nM) for assay validation for 50 minutes at 37 °C. Unstimulated cells served as negative controls. Reactions were arrested on ice, and cells were stained with fluorochrome-conjugated monoclonal antibodies targeting CD123, HLA-DR, CD63, and CD203c for 30 minutes at 4 °C. Basophil populations were gated as CD123+/HLA-DR-events, and functional activation was quantified by the percentage of CD63+ cells and the median fluorescence intensity of CD203c.

### Isolation and Culture of Feline Fetal Fibroblasts

Feline fetal tissues were obtained from domestic cats (*Felis catus*) during routine spay procedures conducted as part of an educational trap-neuter-return (TNR) programme at Seoul National University. The foetuses, measuring approximately 2.5 cm in length, were collected under the clinical supervision of Prof. Goo Jang (DVM, PhD). All clinical interventions, including anaesthesia and surgical procedures on donor queens, were performed exclusively by the veterinary staff of the Seoul National University Veterinary Medical Teaching Hospital (SNU VMTH). Researchers were provided with euthanised foetuses and were not involved in any live animal handling.

Primary feline foetal fibroblasts were derived from the dorsal skin of the foetuses following an established protocol for embryonic fibroblast isolation^38^. Briefly, Skin tissues were minced and enzymatically dissociated using 10 mg/ml Collagenase I at 37°C overnight. The resulting cell suspension was processed through a 70-µm strainer and collected via centrifugation at 700×g for 5 minutes. Cells were maintained in Dulbecco’s modified Eagle’s medium (DMEM; Cat# 11995073) supplemented with 10% foetal bovine serum (FBS; Cat# 12483020) and 10 U/ml penicillin-streptomycin. The cultures were incubated at 37 °C in a humidified environment with 5% CO₂. Feline foetal fibroblasts at passage 3 were used for all subsequent experiments.

### CRISPR/Cas9-mediated genome editing and proliferation monitoring

CRISPR/Cas9-mediated genome editing of feline foetal fibroblasts was executed using a ribonucleoprotein (RNP) complex paired with an antisense single-stranded oligodeoxynucleotide (ssODN) donor template. A single guide RNA (sgRNA, 5’-CACTTGCTCAACATATTCGTCGG-3’) targeting the Fel d 1 locus was designed utilising CRISPR RGEN Tools. The RNP complex was formulated by combining 600 ng/µl Cas9 protein (Invitrogen) with 300 ng/µl sgRNA. The antisense ssODN donor (left homology arm: 107 bp, right homology arm: 90 bp) was engineered to harbour the K29G substitution alongside a silent mutation to prevent Cas9 re-cleavage.

Transfection of approximately 2×10⁵ cells per reaction was performed using the Neon™ NxT Electroporation System (Thermo Fisher Scientific) in the presence of 1.8 µM Cas9 Electroporation enhancer and 70 µM HDR enhancer (IDT, 10007910). Following transfection, cells were subjected to limiting dilution to initiate single-cell culture. Emerging clones were sequentially expanded through a stepwise scale-up process into progressively larger culture vessels. Upon reaching approximately 80% confluency in 35-mm dishes, cells from each clone were harvested for genomic DNA (gDNA) extraction. The target locus was PCR-amplified, and successful integration of the targeted K29G substitution was verified via Sanger sequencing.

Cellular viability and growth kinetics of the edited fibroblasts were assessed by longitudinally monitoring the population doubling time (PDT). Wild-type and K29G-edited cell lines were continuously cultured under standard conditions, and the PDT was calculated at each passage from passage 8 through passage 19. Cells were counted at each split using a haemocytometer to ensure consistent seeding densities, and doubling times were derived to evaluate the fundamental cellular fitness following the gene-editing intervention.

## Acknowledgements

This work was supported by the Korea Health Technology R&D Project through the Korea Health Industry Development Institute (KHIDI), funded by the Ministry of Health & Welfare, Republic of Korea (RS-2025-25459531 to Y.C.); the Medical Research Center for Innovative Control of Cardiovascular Remodeling Diseases (RS-2025-02213506 to Y.C.); National Research Foundation of Korea (NRF) grants funded by the Korean government (MSIT) (2020R1A5A2031185 to Y.C. and 2710004423 to S.-Y.Y.); the National Research Foundation Singapore (NRF-MSG-2023-0003 to J.K.L.); the National University of Singapore (NUHSRO/2024/064/NUSMed/05/SynCTI2.0 to J.K.L.); and a Start-up Grant (7100103 to J.K.L.).

