## Supplementary for "Structure-guided deimmunisation of Fel d 1 suppresses allergic effector functions ex vivo"

**Table S1. Sequences of natural and recombinant Fel d 1.** Full-length sequences of native Fel d 1 (chain 1 and chain 2 separately) and the recombinant single-chain fusion construct used in this study. Mutated positions are indicated in bold and underlined.

| | | Position | Sequence |
|---|---|---|---|
| nFel d 1 | Chain 1 | <br>1<br>21<br>41<br>61 | **12345678901234567890**<br>EICPAVKRDVDLFLTGTPDE<br>YVEQVAQYKALPVVLENARI<br>LKNCVDAKMTEEDKENALSV<br>LDKIYTSPLC |
| | Chain 2 | <br>1<br>21<br>41<br>61<br>81 | **12345678901234567890**<br>VKMAETCPIFYDVFFAVANG<br>NELLLDLSLTKVNATEPERT<br>AMKKIQDCYVENGLISRVLD<br>GLVMTTISSSKDCMGEAVQN<br>TVEDLKLNTLGR |
| rFel d 1 | | <br>1<br>21<br>41<br>61<br>81<br>101<br>121<br>141<br>161 | **12345678901234567890**<br>EICPAVKRDVDLFLTGTPDE<br>YVEQVA**Q**Y**KALP**VVLENARI<br>LKNCVDAKMTEEDKENALSL<br>LDKIYTSPLCVKMAETCPIF<br>YDVFFAVANG**NELL**LDLSLT<br>KVNATEPERTAMKKIQDCYV<br>ENGLISRVLDGLVMTTISSS<br>KDCMGEAVQNTVEDLKLNTL<br>GR |

**Table S2. Known T- and B-cell epitopes of Fel d 1.**

| **T-cell epitopes** | | | | |
|---|---|---|---|---|
| Chain | Epitope group | HLA type | Range (rFel d 1 range) | Epitope |
| 1 | 1 | DRB5*01:01 | 19-31 | DEYVEQVAQYKAL |
| | | DRB1*01:01 | 25-44 | VAQYKALPVVLENARILKNC |
| | | DRB1*13:01 | 31-43 | LPVVLENARILKN |
| | 2 | DRB1*14:01 | 55-67 | ENALSLLDKIYTS |
| | | DRB1*11:01 | 58-67 | LSLLDKIYTS |
| 2 | 3 | DRB1*04:01 | 22-31 (92-101) | ELLLDLSLTK |
| **B-cell epitopes** | | | | |
| Chain | Epitope group | | Range (rFel d 1 range) | Epitope |
| 1 | 1 | | 25-38 | VAQYKALPVVLENA |
| | 2 | | 46-59 | DAKMTEEDKRNALS |
| 2 | 3 | | 12-25 (82-95) | DVFFAVANGNELLL |

**Table S3. Comprehensive IgE-binding profiles of Fel d 1 variants.** The compiled ELISA results incorporate the initial screening data derived from the highly reactive donor P5 (+) and the non-allergic control N7 (-). Serum from the negative control N7 was utilised twice.

| Variant | N1(-) | N7(-) | N7(-) | N8(-) | P1(+) | P2(+) | P3(+) | P5(+) | Average (-) | Average (+) |
|---|---|---|---|---|---|---|---|---|---|---|
| Q27C | 0.571 | 0.532 | 0.081 | 0.113 | 0.058 | 0.122 | 0.086 | 0.669 | 0.324 | 0.234 |
| K29G | 0.087 | 0.616 | 0.232 | 0.057 | 0.097 | 0.116 | 0.741 | 0.394 | 0.248 | 0.337 |
| A30D | 0.085 | 0.553 | 0.235 | 0.058 | 0.098 | 0.118 | 0.711 | 0.426 | 0.233 | 0.338 |
| A30W | 0.089 | 0.557 | 0.237 | 0.057 | 0.094 | 0.117 | 0.737 | 0.521 | 0.235 | 0.367 |
| L31E | 0.091 | 0.418 | 0.213 | 0.055 | 0.291 | 1.760 | 1.804 | 0.393 | 0.194 | 1.062 |
| P32G | 0.081 | 0.597 | 0.179 | 0.056 | 0.007 | 0.112 | 0.666 | 0.399 | 0.228 | 0.296 |
| N91D | 0.080 | 0.436 | 0.177 | 0.057 | 0.098 | 0.112 | 0.715 | 0.426 | 0.187 | 0.338 |
| E92D | 0.091 | 0.368 | 0.226 | 0.060 | 0.098 | 0.134 | 0.774 | 0.373 | 0.186 | 0.345 |
| L93E | 0.092 | 0.705 | 0.217 | 0.062 | 0.012 | 0.119 | 0.759 | 0.587 | 0.269 | 0.369 |
| L93N | 0.089 | 0.560 | 0.190 | 0.057 | 0.056 | 0.120 | 0.752 | 0.412 | 0.224 | 0.335 |
| L93Q | 0.084 | 0.571 | 0.211 | 0.056 | 0.091 | 0.123 | 0.715 | 0.443 | 0.230 | 0.343 |
| L94N | 0.083 | 0.353 | 0.208 | 0.056 | 0.090 | 0.124 | 0.726 | 0.363 | 0.175 | 0.326 |
| L94P | 0.079 | 0.419 | 0.159 | 0.057 | 0.088 | 0.116 | 0.703 | 0.440 | 0.179 | 0.337 |
| rFel d 1 | 0.081 | 0.554 | 0.166 | 0.061 | 0.334 | 1.498 | 1.298 | 1.523 | 0.215 | 1.163 |
| nFel d 1 | 0.068 | 0.477 | 0.187 | 0.054 | 0.422 | 2.158 | 2.262 | 1.312 | 0.197 | 1.539 |
| Negative | 0.053 | 0.488 | 0.187 | 0.058 | 0.090 | 0.052 | 0.701 | 0.455 | 0.197 | 0.324 |

**Figure S1. SDS-PAGE analysis of recombinant wild-type and variant Fel d 1 proteins.** Wild-type rFel d 1 and nFel d 1 show correct molecular weights (~18-20 kDa) and comparable purity following refolding and purification.

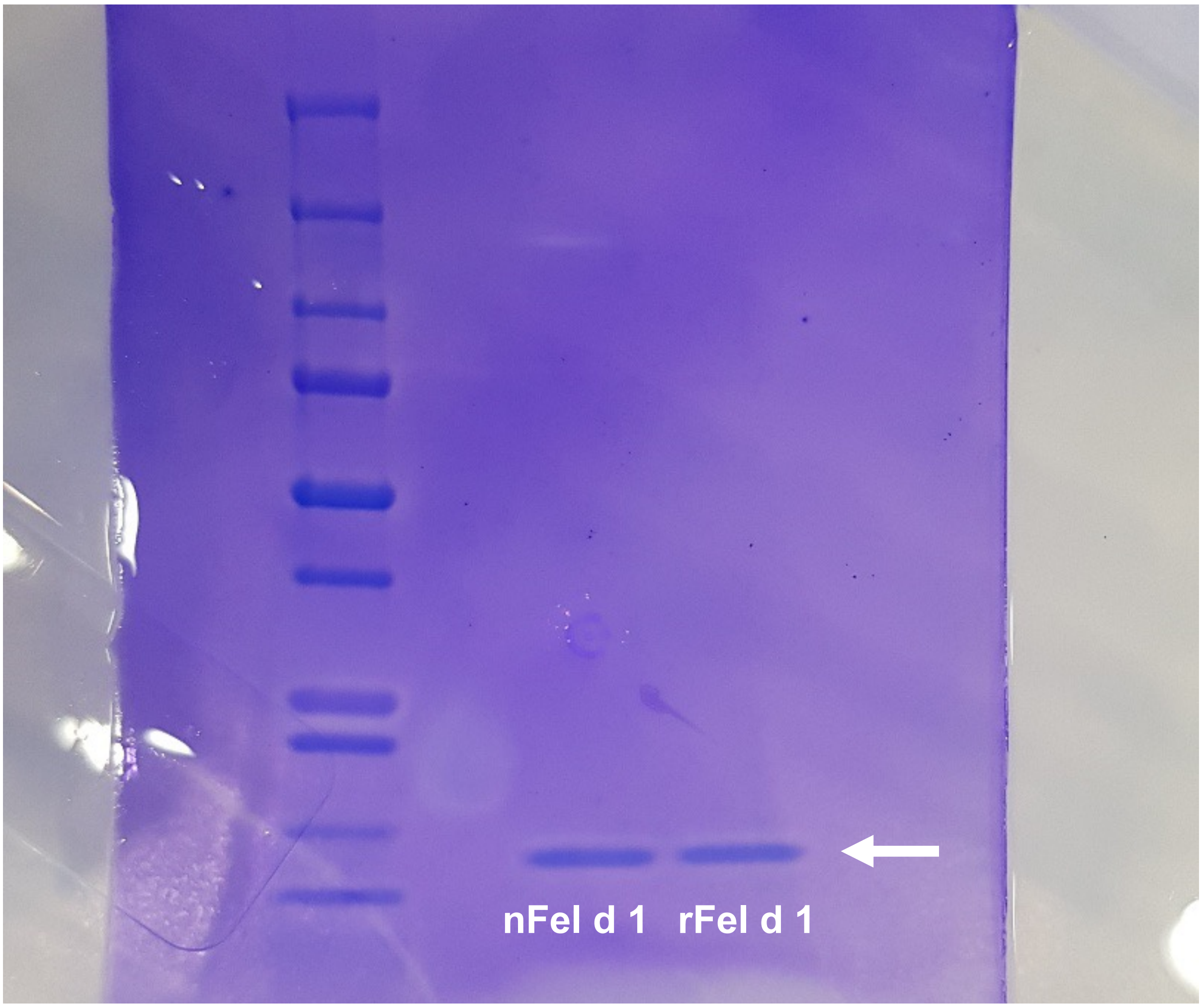

**Figure S2. Validation of basophil activation assay using fMLP.** Flow cytometry plots from donor P5 show a dose-dependent increase in CD123$^{+}$ CD63$^{+}$ basophil population upon fMLP stimulation.

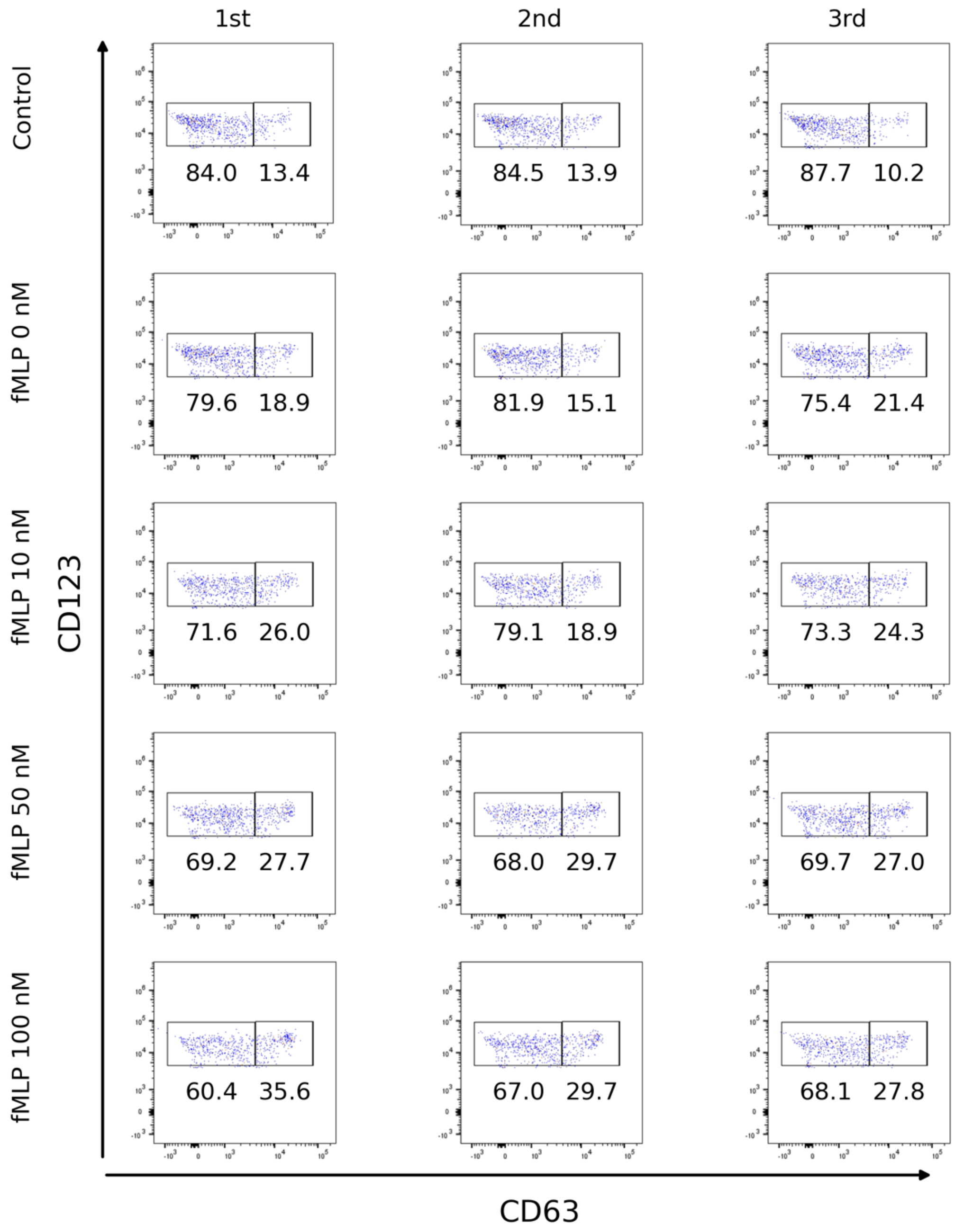

**Figure S3. Functional basophil responses to rFel d 1 and variant K29G.** FACS profiles from donors P2, P3, and P8 demonstrate robust activation by rFel d 1 and minimal activation by K29G.

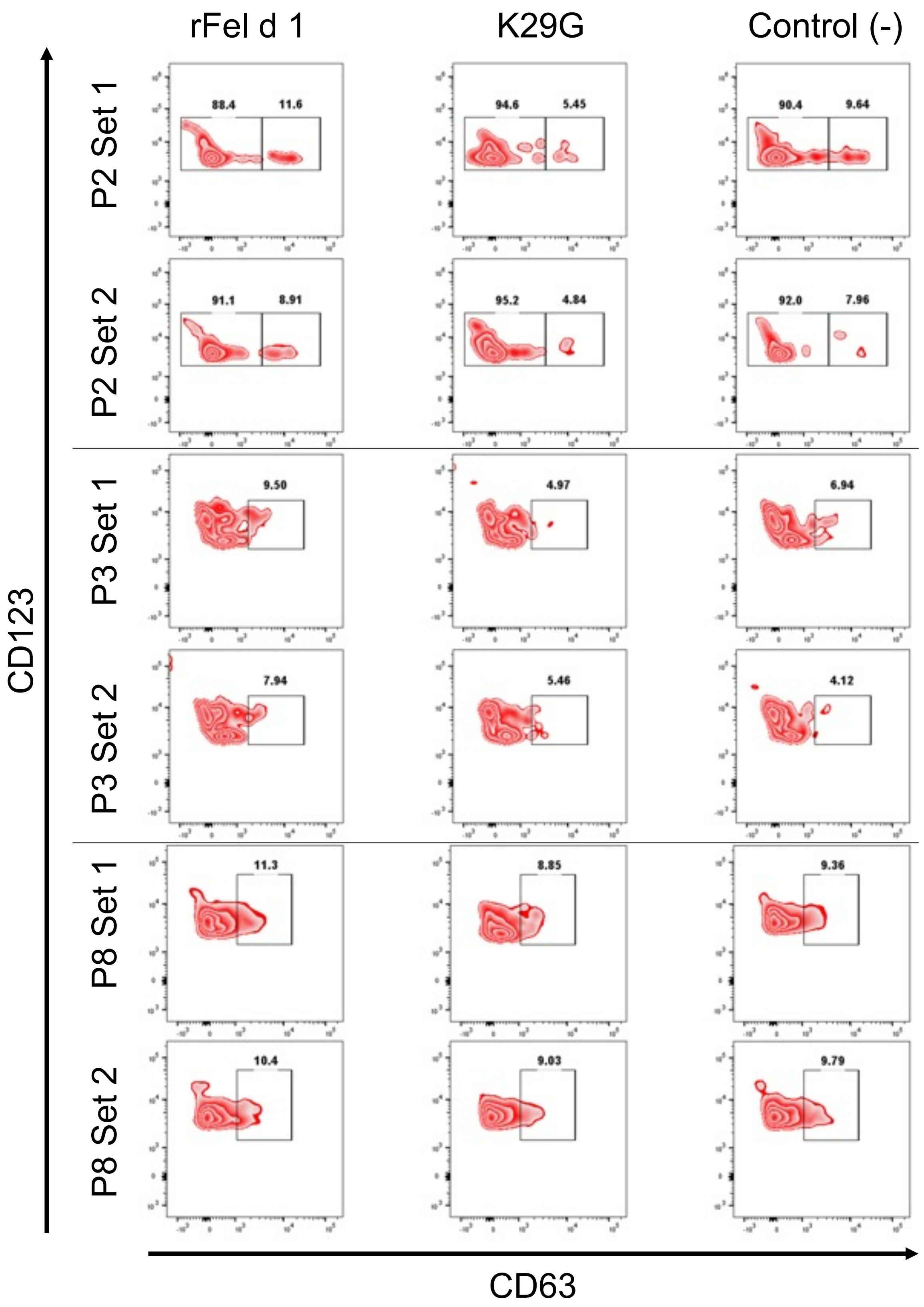

**Figure S4. Genetic confirmation of K29G-edited feline fibroblasts.** Sequence alignment displaying the wild-type reference, the single-stranded oligodeoxynucleotide (ssODN) donor template, and the resulting K29G-edited cell line. The protospacer adjacent motif (PAM, green line) was disrupted by a silent mutation (C→A, green highlight) in the ssODN, preventing Cas9 re-cleavage while preserving the native proline (P) residue. The targeted K29G substitution (AAA→GGA, red boxes) was successfully integrated, as confirmed by Sanger sequencing chromatograms of the established clonal line.

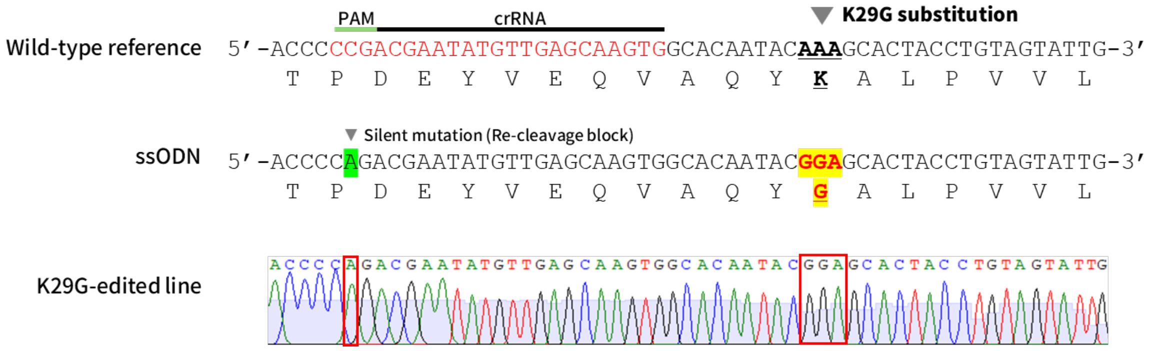
